# The transcriptome of soybean reproductive tissues subjected to water deficit, heat stress, and a combination of water deficit and heat stress

**DOI:** 10.1101/2023.01.29.526088

**Authors:** Ranjita Sinha, Sai Preethi Induri, María Ángeles Peláez-Vico, Adama Tukuli, Benjamin Shostak, Sara I. Zandalinas, Trupti Joshi, Felix B. Fritschi, Ron Mittler

## Abstract

Global warming and climate change are driving an alarming increase in the frequency and intensity of extreme climate events, such as droughts, heat waves, and their combination, inflicting heavy losses to agricultural production. Recent studies revealed that the transcriptomic responses of different crops to water deficit (WD) or heat stress (HS) is very different from that to a combination of WD+HS. In addition, it was found that the effects of WD, HS, and WD+HS are significantly more devastating when these stresses occur during the reproductive growth phase of crops, compared to vegetative growth. As the molecular responses of different reproductive and vegetative tissues of plants to WD, HS, or WD+HS could be different from each other, and these differences could impact many current and future breeding and/or engineering attempts to enhance the resilience of crops to climate change, we conducted a transcriptomic analysis of different soybean (*Glycine max*) tissues to WD, HS, and WD+HS. Here we present a reference transcriptomic dataset that includes the response of soybean leaf, pod, anther, stigma, ovary, and sepal to WD, HS, and WD+HS conditions. Mining this data set for the expression pattern of different stress-response transcripts revealed that each tissue had a unique transcriptomic response to each of the different stress conditions. This finding is important as it suggests that attempting to enhance the overall resilience of crops to climate change could require a coordinated approach that simultaneously alters the expression of different groups of transcripts in different tissues in a stress-specific manner.

**SIGNIFICANCE STATEMENT:** A reference transcriptomic dataset of different reproductive tissues of soybean subjected to water deficit, heat stress, and their combination, generated by this study, reveals that different tissues display different responses to these stress conditions. Attempting to enhance the resilience of crops to different stress combinations, associated with climate change, might therefore require simultaneously altering the expression of different sets of transcripts in different tissues in a coordinated and stress-specific manner.

## INTRODUCTION

Global warming is driving an alarming increase in the frequency and intensity of climate extremes, such as droughts, floods, cold snaps and/or heat waves, inflicting heavy losses to agricultural production and causing food insecurities that destabilize different societies worldwide (Lobell and Gourdji, 2012; Lesk et al., 2016; Alizadeh et al., 2020; Overpeck and Udall, 2020; Brás et al., 2021; Masson-Delmotte et al., 2021; Zandalinas et al., 2021; Lesk et al., 2022). In addition to the costly effects of each of these individual climate events (*e*.*g*., droughts, floods, or heat waves), in some instances two or more climate extremes can impact crops simultaneously, *e*.*g*., when a heat wave occurs during droughts or floods (*e*.*g*., Mazdiyasni and AghaKouchak, 2015; Hao et al., 2018; Alizadeh et al., 2020; Overpeck and Udall, 2020; Zandalinas et al., 2021; Lesk et al., 2022). Historically, a combination of drought and heat wave had a devastating impact on agriculture, surpassing the effects of each individual condition and resulting in heavy losses to grain production in major crops such as maize, soybean, rice, and wheat (Mittler, 2006; Lobell and Gourdji, 2012; Suzuki et al., 2014; Mahrookashani et al., 2017; Cohen et al., 2021b; Bheemanahalli et al., 2022; Liu et al., 2022). Multiple studies have now revealed that the occurrence of drought and heat wave episodes is gradually increasing in recent years due to global warming and climate change, posing a heightened risk to agriculture (*e*.*g*., AghaKouchak et al., 2014; Alizadeh et al., 2020; Potopová et al., 2021; Zandalinas et al., 2021). The effects of water deficit and heat stress combination on crop yield was also found to be significantly more devastating when the two stresses coincide during the reproductive growth phase of crops, compared to during vegetative growth (*e*.*g*., Suzuki et al., 2014; Mahrookashani et al., 2017; Lawas et al., 2018; Cohen et al., 2021b; Sinha et al., 2021; Bheemanahalli et al., 2022; Liu et al., 2022).

The impact of water deficit, heat stress, and/or their combination on reproductive processes of major crops has been the subject of intense research efforts and improving the tolerance of reproductive tissues to these stress conditions is a major goal of breeders and the agricultural biotech industry worldwide (*e*.*g*., Prasad et al., 2008; Fang et al., 2010; Rang et al., 2011; Jin et al., 2013; Su et al., 2013; Jagadish et al., 2015; Oury et al., 2016a; Mahrookashani et al., 2017; Lawas et al., 2018; Bheemanahalli et al., 2019; Gaur et al., 2019; Lippmann et al., 2019; Reichardt et al., 2020; Da Costa et al., 2022; Ishimaru et al., 2022; Rivero et al., 2022). As the development and maturation of reproductive tissues, the fertilization process, embryogenesis, and seed maturation, involve the coordination of multiple developmental, stress-response, and/or programmed cell death (PCD) pathways, under controlled conditions, it was hypothesized that any stress, such as heat stress (HS), water deficit (WD), or their combination (*i*.*e*., WD+HS), could alter the balance between development and stress response pathways and impair the entire reproductive process (Endo et al., 2009; Jin et al., 2013; Su et al., 2013; Ma et al., 2014; Oury et al., 2016b; Djanaguiraman et al., 2018; Begcy et al., 2019; Bheemanahalli et al., 2019; Fábián et al., 2019; del Olmo et al., 2019; Santiago and Sharkey, 2019; Hedhly et al., 2020; Lohani et al., 2020; Chaturvedi et al., 2021; Sze et al., 2021; Santiago et al., 2021; Sinha et al., 2021; Zhang et al., 2021; Lu et al., 2022; Mareri and Cai, 2022). For example, the activation of stress response pathways, such as those inducing desiccation tolerance, could occur too early during the pollen or seed maturation processes under WD conditions, and the activation of PCD pathways in the tapetum during the development of pollens could occur too early under HS conditions. Of particular importance to the synchronization of reproductive processes under controlled growth conditions are transient changes in the levels of stress hormones such as abscisic acid (ABA) and jasmonic acid (JA), and/or reactive oxygen species (ROS), that accompany baseline reproductive processes and play a key role in harmonizing them. The levels of ABA, JA, and/or ROS can however be altered in reproductive tissues during stresses such as WD, HS, or WD+HS, which may disrupt the delicate and coordinated function of these signaling molecules, required for the baseline regulation of plant reproductive processes, and impair overall reproduction and yield (*e*.*g*., reviewed in Sze et al., 2021; Santiago et al., 2021; Sinha et al., 2021).

We recently conducted a comparative transcriptomic analysis of the response of whole flower, pod, and leaf, to WD, HS, and WD+HS, in soybean (*Glycine max*) plants and found that the transcriptomic response of reproductive tissues (whole flower or pod) was different from that of leaf to each of these stresses, as well as their combination (Cohen et al., 2021a; Sinha et al., 2022b; Sinha et al., 2022a). In addition, we found that, in contrast to stomata on leaf which remained closed during WD+HS combination to prevent water loss, stomata of sepal and pod remained open during this stress combination (Sinha et al., 2022a; Sinha et al., 2022b). This differential stomatal response between reproductive (flower and pod) and vegetative (leaf) tissues was accompanied by a differential transpiration response (high in flower and pod and low in leaf), that protected reproductive tissues from overheating during the stress combination (at the expense of leaves). We termed this newly discovered acclimation response of plants ‘Differential transpiration’ (Sinha et al., 2022b).

As the transcriptomic response of different tissues of plants to WD+HS combination was found to be different between different tissues (whole flower, pod, and leaf), and this difference could impact many current and future breeding and/or engineering attempts to enhance the resilience of crops to climate change, we expanded the transcriptomic analysis of different plant tissues to WD+HS combination to include anther, stigma, ovary, and sepal. Here we present a reference transcriptomic dataset that includes the response of soybean leaf, pod, anther, stigma, ovary, and sepal to WD, HS, or a combination of WD+HS. Mining this data set for the expression pattern of HS-, WD-, hormone-, and ROS-related transcripts revealed distinct expression patterns for specific pathways in different tissues during different stress conditions and increased our overall understanding of the different molecular processes that occur in plants during a combination of WD+HS. Future mining of the dataset presented in this study could lead to the identification of new pathways and genes that may be used to enhance the resilience of crops to heat waves, droughts, and their combination, preventing losses estimated in billions of dollars to agriculture and increasing food security worldwide. Our findings are also important as they suggest that attempting to enhance the overall resilience of crops to different stresses, and/or their combinations, could require strategies that simultaneously target and coordinate multiple stress- response pathways in different tissues.

## RESULTS AND DISCUSSION

### Growth conditions, sampling, and differences in the basal transcriptome of different tissues

To study the transcriptomic response of different soybean (*Glycine max, cv Magellan*) tissues to conditions of WD, HS, or a combination of WD+HS, we grew plants under controlled growth conditions in chambers as previously described (Cohen et al., 2021a; Sinha et al., 2022a; Sinha et al., 2022b). When plants started flowering (R1 stage) we induced conditions of WD, HS, or WD+HS (Sinha et al., 2022b) and maintained these conditions for 10-15 days before starting to sample the different plant tissues. This design ensured that all sampled tissues developed on plants subjected to the different stress conditions. In addition, and as previously described, all flowers used for the transcriptomic analysis presented in the current study were at stages II and III (unopen flowers undergoing self-pollination) from plants at the R2 stage (Sinha et al., 2022b), and all pods were at a length of about 3 cm and contained developing seeds (Sinha et al., 2022a). Leaves and pods were sampled and immediately flash frozen in liquid nitrogen as previously described (Sinha et al., 2022a; Sinha et al., 2022b; Cohen et al., 2021a), while flowers were dissected as shown in Figure 1a and the different flower tissues (sepals, anthers, ovary, and stigma) were immediately flash frozen in liquid nitrogen. All tissues were processed for RNA-Seq analysis using the same protocols and all RNA was sequenced by Novogene co. Ltd (https://en.novogene.com/; Sacramento, CA, USA) using NovaSeq 6000 PE150 [Supplementary Tables S1-S36; GSE218146, GSE213479, GSE153951, and GSE186317, generated by this study and Sinha et al., (2022a), Sinha et al., (2022b), and Cohen et al., (2021a)]. All previously- (leaf and pod) and newly- (sepals, anthers, ovary, and stigma) generated raw RNA-Seq data were re-processed together and subjected to all post sequencing analyses steps as described in Sinha et al., (2022b).

**Figure 1.**
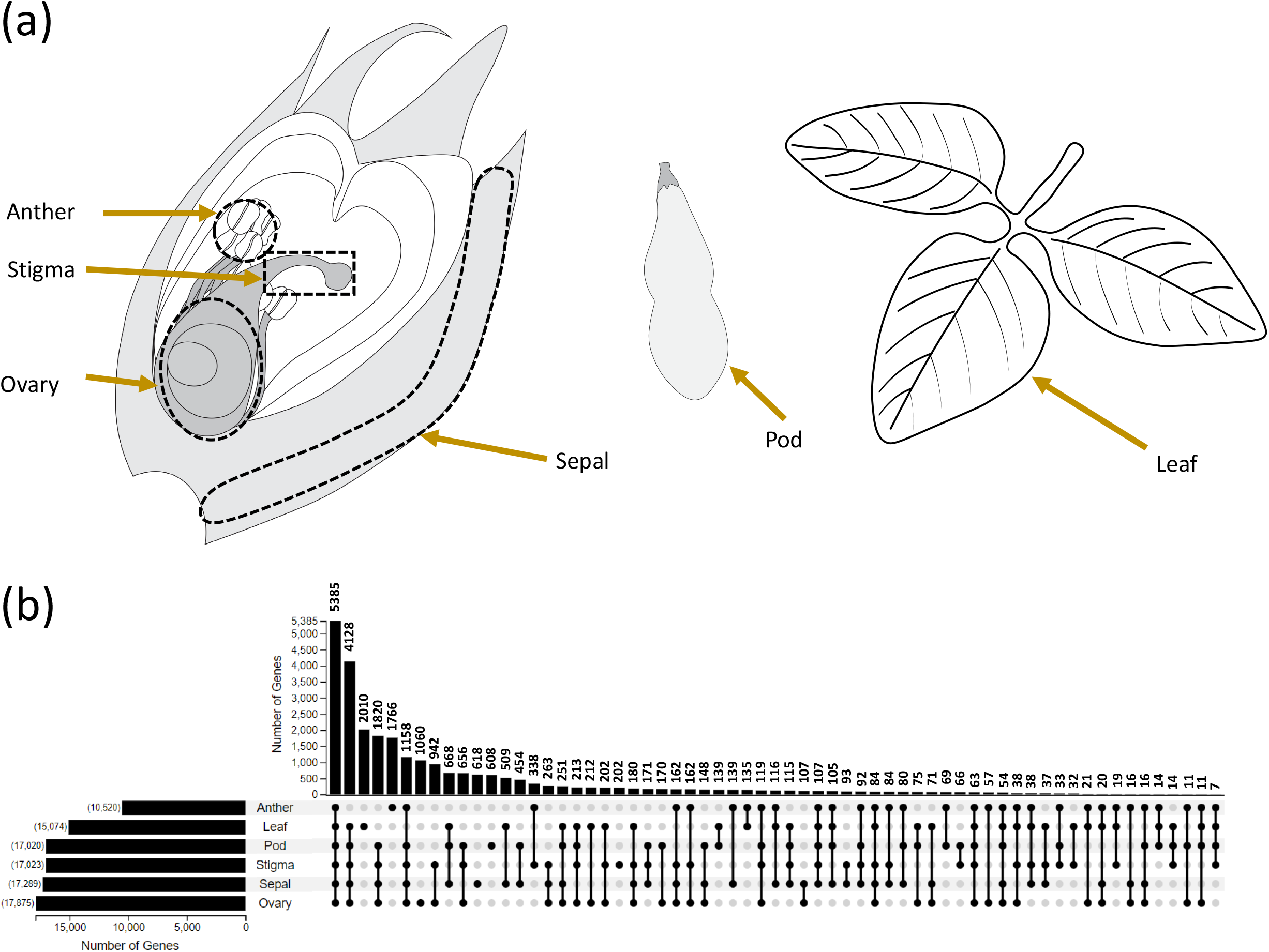
Similarities and differences in the basal transcriptome of the different soybean tissues obtained from plants grown under controlled growth conditions. (a) An illustration depicting the different plant organs used for the transcriptome analysis. (b) An UpSet plot of the overlap in basal transcript expression between the different tissues obtained from plants grown under control conditions. All experiments were conducted in 3 biological repeats each with tissues pooled from 8-20 different plants (depending on tissue type).

To initiate our stress-response RNA-Seq tissue specific analysis in soybean, we compared the basal expression level of all transcripts with an FPKM (Fragments Per Kilo base of transcript per Million mapped fragments) value higher than 5 under controlled growth conditions, between the different tissues (Supplementary Tables S1-S6). As shown in Figure 1b, leaf, anther, ovary, sepal, pod, and stigma had more than 2,000, 1,700, 1,000, 600, 600, and 200 unique transcripts expressed, respectively (Supplementary Tables S7-S12). In contrast, more than 5,300 transcripts were commonly expressed in all tissues, while over 1,100 transcripts were common to all reproductive tissues, and over 940 transcripts were common to stigma and ovary (Figure 1b; Supplementary Tables S13-S15). The analysis shown in Figure 1 demonstrates that different tissues contain different sets of transcripts, and that stigma may be different from ovary by a few hundred transcripts only. Taken together, the sets of transcripts obtained from each tissue (Figure 1b; Supplementary Tables S1-S33) indicate that the sampling strategy used (Figure 1a) could discern differences in the response of each tissue to WD, HS, or WD+HS, and that comparisons performed between these tissues could reveal biologically significant differences in the response of each tissue to the different stress treatments studied.

### Differential transcriptomic responses to WD, HS, or a combination of WD+HS within and between the different tissues

To examine the response of each individual tissue to WD, HS, or WD+HS and compare responses among tissues, we generated Venn diagrams for each tissue (Figure 2). This analysis revealed that each tissue displayed a transcriptomic response that was specific to WD+HS. In anther and pod this response contained over 7,500 and 9,500 transcripts respectively, while in leaf, sepal, ovary, and stigma it contained over 4,500, 3,600, 5,000, and 4,900 transcripts, respectively. This finding suggests that compared to all other tissues, anther and pod may require a more extensive response to the WD+HS combination. An additional interesting finding that emerged from the data shown in Figure 2 is that the transcriptomic response of all reproductive tissues studied included fewer transcripts (359, 1155, 204, 525, and 822 transcripts in pod, sepal, anther, stigma, and ovary, respectively) than that of leaf (over 4,600 transcripts) in response to WD stress. This finding suggests that in contrast to leaf, reproductive tissues might be better protected during WD stress, perhaps because they represent a primary sink tissue of the plant (Harrison et al., 2022). In contrast to the transcriptomic responses of leaf, pod, and sepal (over 3,000, 2,400, and 4,300 transcripts, respectively), the transcriptomic responses of anther, stigma, and ovary to HS was more robust (over 7,000 transcripts in each tissue). This finding could suggest that anther, stigma, and ovary are more sensitive to HS and require a more extensive transcriptomic response to acclimate to these conditions than leaf, pod, and sepal.

**Figure 2.**
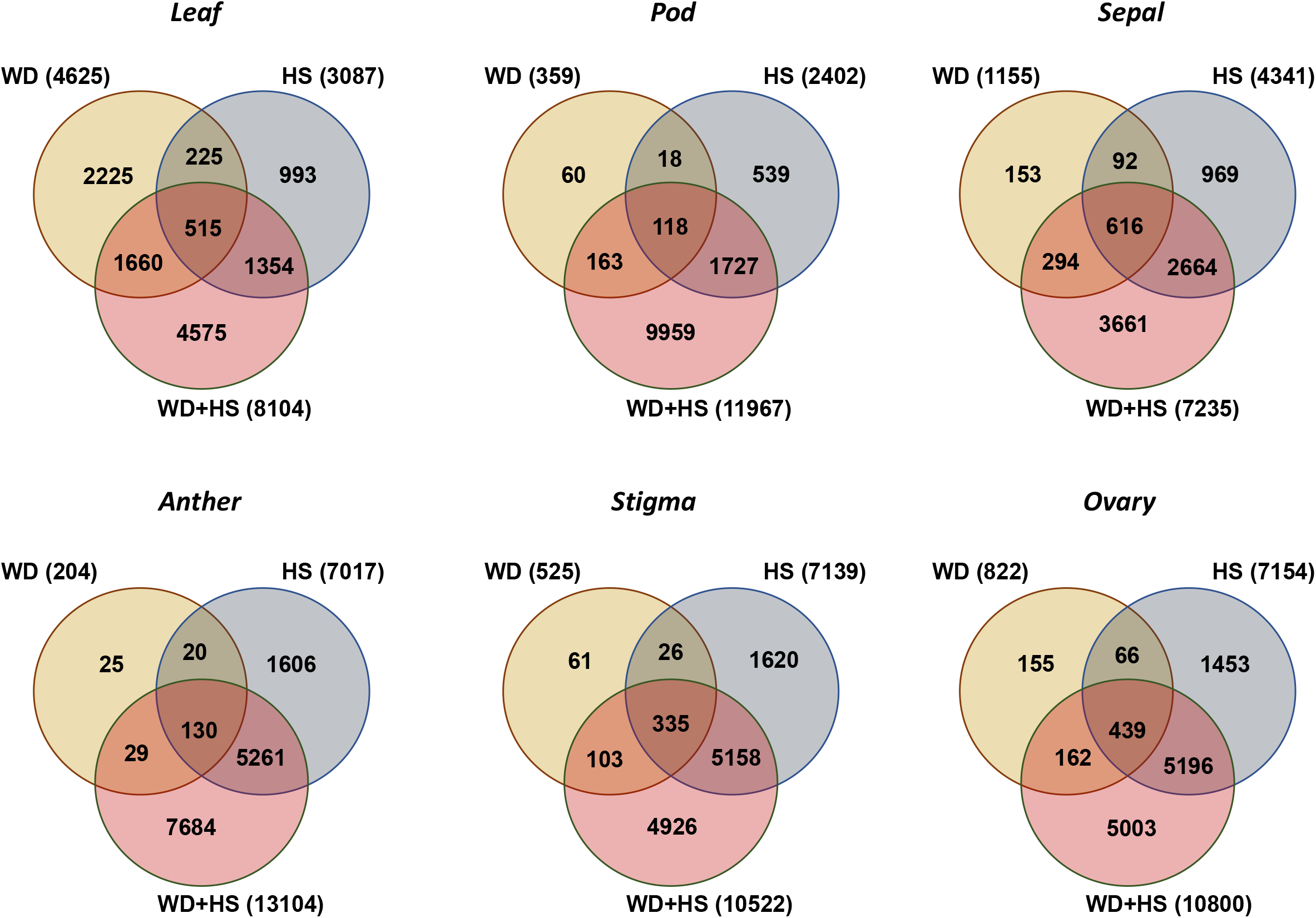
Differential transcriptomic responses to water deficit, heat stress, or a combination of water deficit and heat stress in each of the different tissues. Venn diagrams of the overlap between the transcriptomic responses of leaf, pod, sepal, anther, stigma, and ovary to water deficit, heat stress, or a combination of water deficit and heat stress are shown. All experiments were conducted in 3 biological repeats each with tissues pooled from 8-20 different plants (depending on tissue type). Significant changes in transcript expression compared to control were defined as adjusted P < 0.05 (negative binomial Wald test followed by Benjamini–Hochberg correction). Abbreviations: WD, water deficit; HS, heat stress, WD+HS, a combination of WD and HS.

To compare between the responses of the different tissues to HS, WD, or WD+HS, we used UpSet plots (Figure 3). This analysis revealed a very low overlap between the total transcriptomic response of each tissue to WD (0 transcripts common to all tissues), with leaf and sepal displaying the highest similarity in their response to WD (Figure 3a). A much higher overlap between the total transcriptomic response of the different tissues was found to HS, with stigma and ovary and stigma and anther showing the greatest overlap in transcript expression to this stress (Figure 3b). Interestingly, only 74 transcripts were common to all tissues in response to HS, demonstrating that each of the different tissues had a unique transcriptomic response to HS (Figure 3b). A unique response was also found in all tissues to the combination of WD+HS with anther, pod, leaf, and ovary showing the highest number of unique WD+HS transcripts (over 4,300, 2,600, 2,100, and 1,300, respectively). In contrast to WD (Figure 3a), and HS (Figure 3b), a good overlap was however found between the total transcriptomic response of each tissue to WD+HS, with ovary and stigma, anther and stigma, and anther and pod, showing the greatest overlap in transcript expression (Figure 3c). In addition, more than 560 transcripts were found to be common to the total transcriptomic response of all tissues to WD+HS (Figure 3c). When comparing the transcriptomic responses that were specific in each tissue to WD+HS (Figure 2), a much lower overall similarity was nevertheless found between the different tissues with only 16 transcripts common to all tissues (Figure 3d). Anther and pod, pod and leaf, and pod and ovary displayed the highest overlap between the different tissues (905, 706, and 650 transcripts, respectively) to WD+HS (Figure 3d).

**Figure 3.**
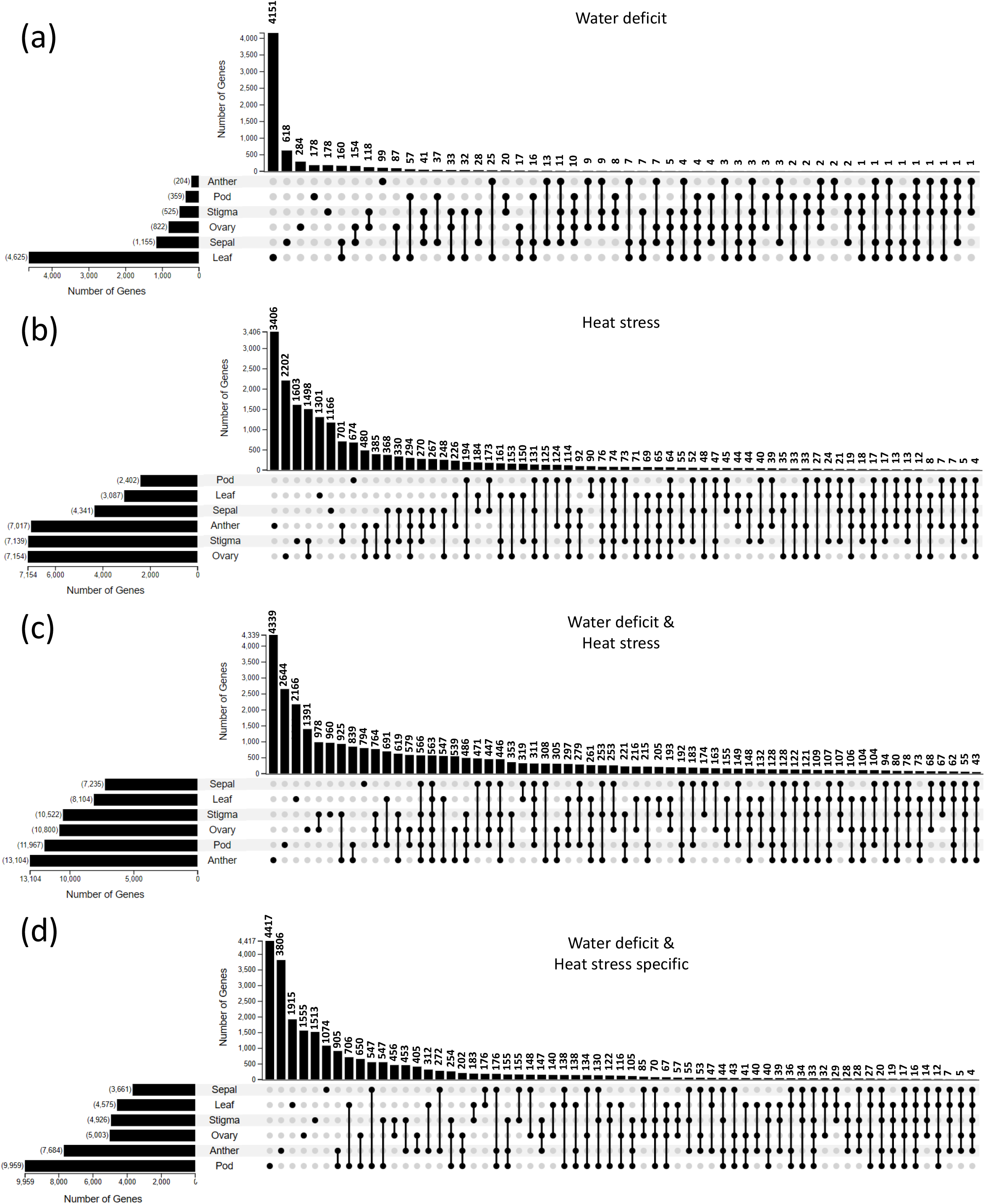
Overlap between the transcriptomic responses of leaf, pod, sepal, anther, stigma, and ovary to water deficit, heat stress, or a combination of water deficit and heat stress. UpSet plots depicting the overlap in transcriptomic responses to water deficit (a), heat stress (b), or a combination of water deficit and heat stress (c) between leaf, pod, sepal, anther, stigma, and ovary are shown. (d) An UpSet plot showing the overlap between the different transcripts that are uniquely expressed in each tissue in response to a combination of water deficit and heat stress (From Figure 2). All experiments were conducted in 3 biological repeats each with tissues pooled from 8-20 different plants (depending on tissue type). Significant changes in transcript expression compared to control were defined as adjusted P < 0.05 (negative binomial Wald test followed by Benjamini–Hochberg correction).

The findings presented in Figures 2 and 3 demonstrate that each tissue has a unique transcriptomic response to WD, HS, or WD+HS and that the responses of the different tissues to the different stress conditions vary. These findings suggest that the chances of attempting to induce resilience of all plant tissues to WD, HS, or WD+HS using alterations in the expression pattern of only one gene, or one pathway, are slim, and that a more focused effort should be made to study tissue specific responses to WD, HS, or WD+HS conditions.

### Expression of HS-, WD-, ABA-, and ROS-related transcripts in the different tissues in response to WD, HS, or a combination of WD+HS

To mine the transcriptomic data obtained from the different tissues developing on plants subjected to CT, WD, HS, or WD+HS, we focused on selected pathways that play important roles in the acclimation of plants to HS, WD, and/or oxidative stress (Schöffl et al., 1998; Frank et al., 2009; Giorno et al., 2010; Liu and Howell, 2010; Suzuki et al., 2011, 2014; Howell, 2013; Sun et al., 2013; Fragkostefanakis et al., 2016; Ohama et al., 2017; Zhang et al., 2017; Zandalinas et al., 2018; Singh et al., 2021; Mittler et al., 2022). As illustrated in Figure 4, comparing the expression pattern of heat shock transcription factors (HSFs; Figure 4a) that regulate many cytosolic, mitochondrial and chloroplastic heat stress responses, and endoplasmic reticulum (ER)-related heat response pathways (the unfolded protein response pathway; UPR; Figure 4b), during WD, HS, or WD+HS, revealed several differences in the way each tissue responded to the different stress conditions. In contrast to leaf, for example, reproductive tissues did not alter the expression of many HS-response transcripts in response to WD. In addition, unlike Arabidopsis, in which the UPR was shown to play a prominent role in protecting reproductive tissues during HS (Howell, 2013; Deng et al., 2016), the expression of many transcripts involved in both the ER (UPR) and HSF pathways was elevated in response to HS or WD+HS in soybean reproductive tissues (Figure 4). This finding suggests that HSFs and the UPR pathway are both involved in the response of reproductive and vegetative soybean tissues to these stresses. Although HSFs displayed an overall similar response to HS and WD+HS among tissues, some differences were found between the expression of specific HSFs in response to these two conditions [*e*.*g*., HSFB2B (GLYMA_01G217400) was mostly expressed during HS in flower organs, and HSFA2 (GLYMA_13G105700) was mostly expressed during WD+HS in pod, stigma, and ovary; Figure 4a]. In addition, some ER (UPR) responses were specific to leaf in response to all stresses, while other ER responses were specific to anther in response to HS or WD+HS, and some were specific to leaf in response to WD (Figure 4b).

**Figure 4.**
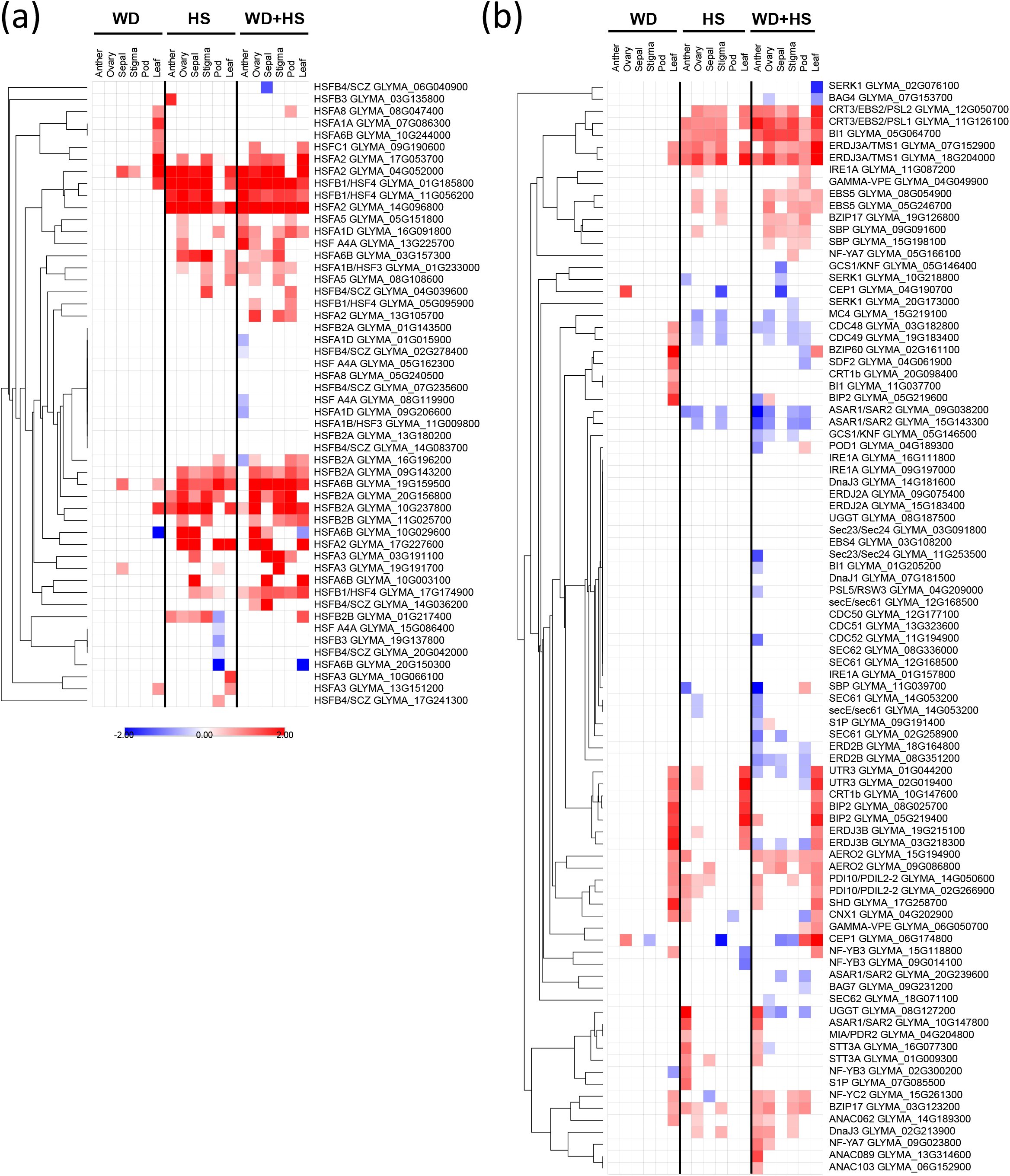
Differential expression of transcripts encoding selected heat stress response transcripts in leaf, pod, sepal, anther, stigma, and ovary in response to water deficit, heat stress, or a combination of water deficit and heat stress. Heat maps depicting the expression of transcripts encoding for soybean heat shock transcription factors (a) and transcripts encoding soybean endoplasmic reticulum unfolded protein response proteins (b) are shown. Only transcripts with a significant expression level compared to control are shown. All experiments were conducted in 3 biological repeats each with tissues pooled from 8-20 different plants (depending on tissue type). Significant changes in transcript expression compared to control were defined as adjusted P < 0.05 (negative binomial Wald test followed by Benjamini–Hochberg correction). Abbreviations: WD, water deficit; HS, heat stress; WD+HS, a combination of WD and HS; HSF, heat shock transcription factor.

Differences in the expression of selected drought-response and ROS scavenging transcripts could also be found between the different tissues in response to the different stress treatments (Figure 5). For example, several drought-response transcripts were highly expressed in pod during WD+HS, but not during WD or HS (Figure 5a). In addition, compared to leaf, several different drought-response transcripts were specifically expressed in reproductive tissues in response to HS, or WD+HS, but not WD (Figure 5a). In contrast to all other tissues, anther displayed an enhanced expression of ROS-scavenging transcripts such as those encoding ascorbate peroxidase 1 and 2 (APX1 and APX2) in response to HS or WD+HS (Figure 5b). In addition, in contrast to leaf, a group of ROS scavenging transcripts including glutathione peroxidase 7 (GPX7) and cupper/zinc superoxide dismutase 3 (CSD3) was upregulated in almost all reproductive tissues in response to HS or WD+HS (Figure 5b).

**Figure 5.**
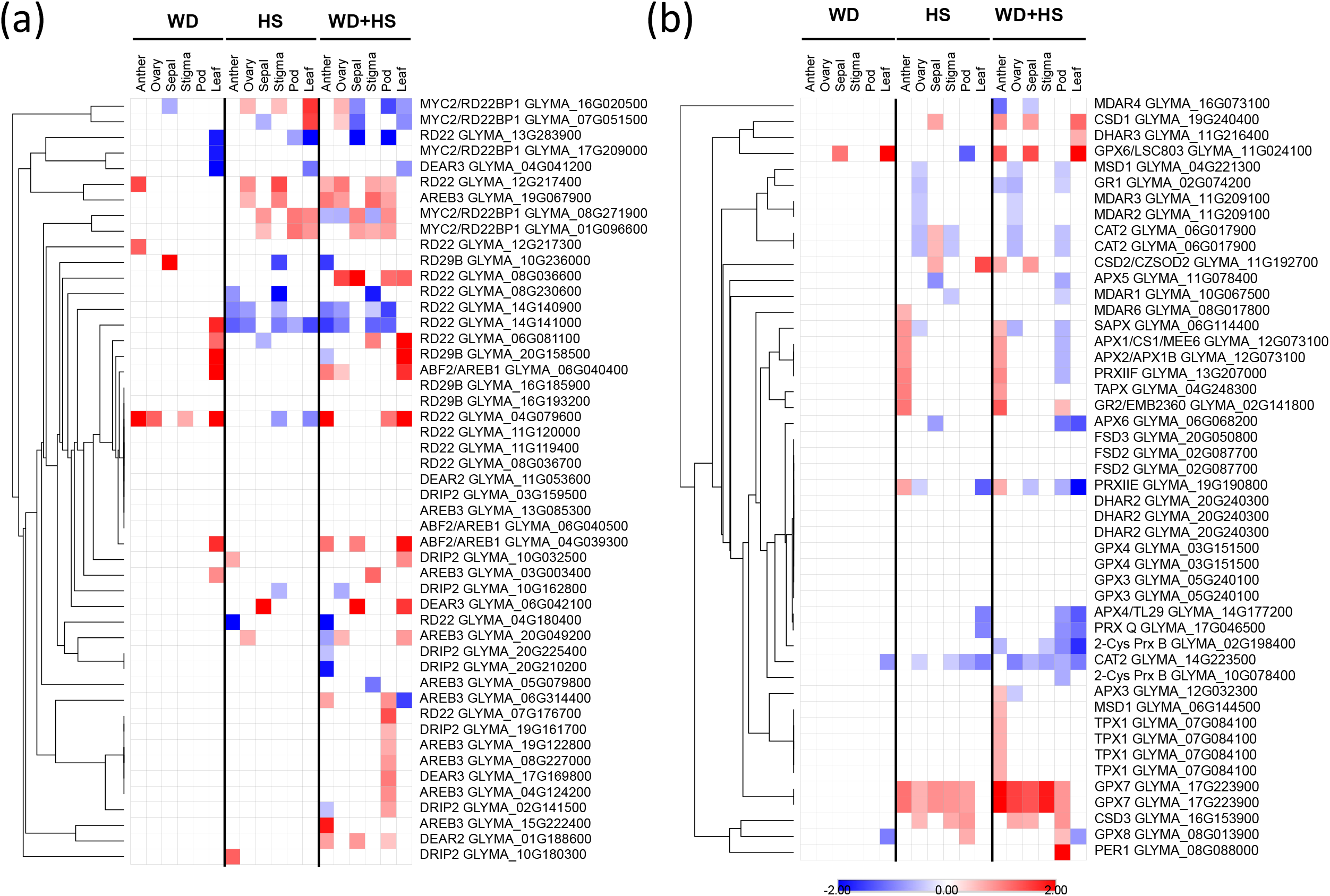
Differential expression of transcripts encoding selected drought and reactive oxygen species response proteins in leaf, pod, sepal, anther, stigma, and ovary in response to water deficit, heat stress, or a combination of water deficit and heat stress. Heat maps depicting the expression of selected soybean drought response transcripts (a) and transcripts encoding different proteins involved in reactive oxygen species scavenging and signaling (b) are shown. Only transcripts with a significant expression level compared to control are shown. All experiments were conducted in 3 biological repeats each with tissues pooled from 8-20 different plants (depending on tissue type). Significant changes in transcript expression compared to control were defined as adjusted P < 0.05 (negative binomial Wald test followed by Benjamini–Hochberg correction). Abbreviations: WD, water deficit; HS, heat stress; WD+HS, a combination of WD and HS.

As shown in Figure 6, while the expression of many transcripts encoding ABA biosynthesis enzymes was upregulated during WD+HS in all tissues, many of these transcripts were not upregulated in response to WD or HS (Figure 6a). In an apparent balancing act, however, the expression of many transcripts encoding ABA degradation enzymes was also upregulated in all tissues during WD+HS (Figure 6a). The expression of one transcript encoding the ABA degradation enzyme ABA 8’-hydroxylase (CYP707A4; GLYMA_07G212700) was specific to sepal, ovary and pod during HS and WD+HS, suggesting that this isozyme of CYP707A might be responsible for the accumulation of dihydrophaseic acid (DPA) during HS and WD+HS in whole flowers and the opening of stomata on sepal and pod during these stress conditions (Sinha et al., 2022b; Sinha et al., 2022a; Figure 6a). Interestingly, compared to leaf, the expression of most transcripts encoding ABA biosynthesis and degradation enzymes was not upregulated in reproductive tissues during WD (Figure 6a). This finding could suggest that during WD, ABA is mobilized from leaves or roots to reproductive tissues, rather than synthesized/degraded in these tissues, or that reproductive tissues are less sensitive than other plant tissues to WD due to being a prime sink tissue (Harrison et al., 2022).

**Figure 6.**
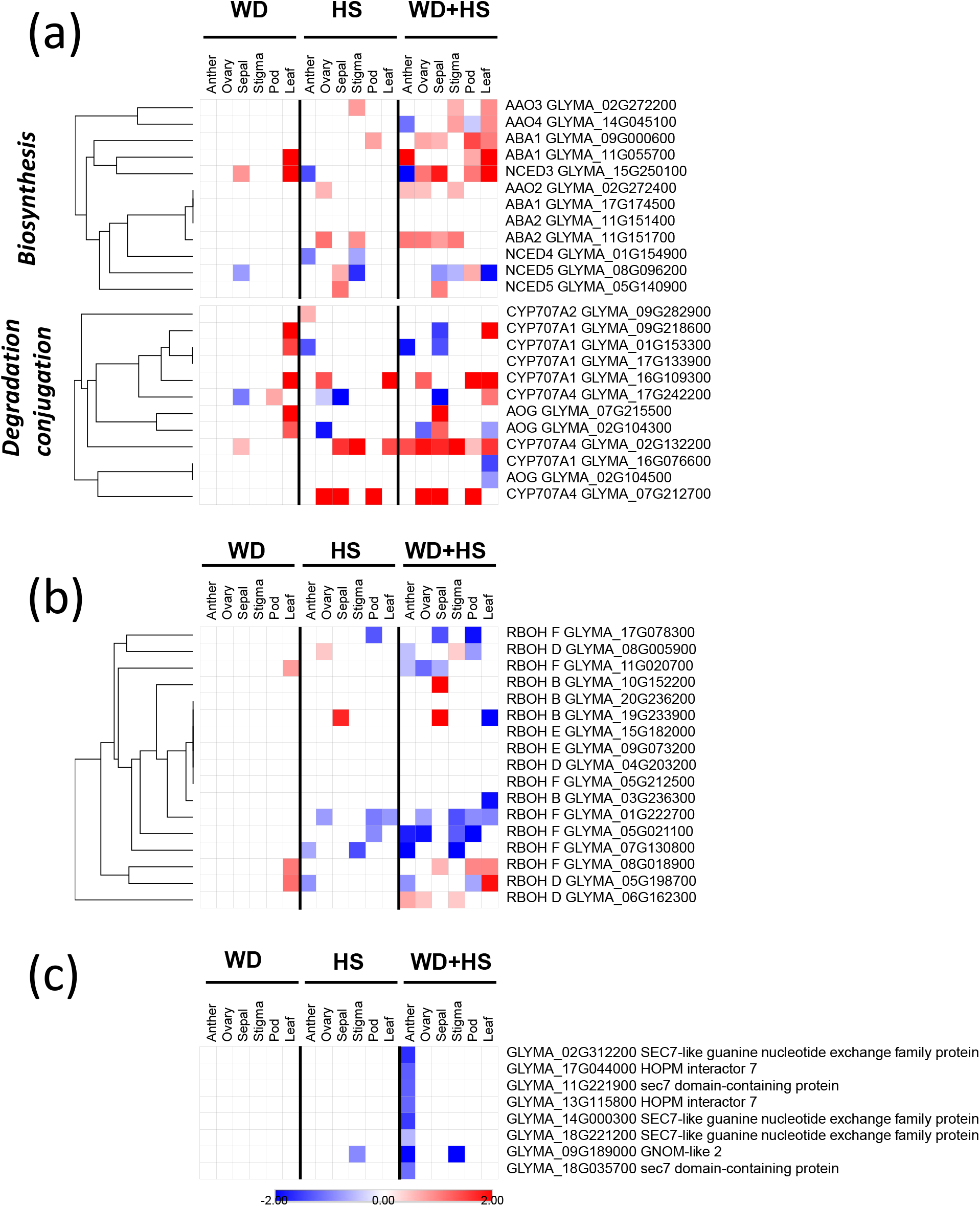
Differential expression of transcripts encoding abscisic acid metabolism, respiratory burst oxidase homologs, and auxin response factor signaling in leaf, pod, sepal, anther, stigma, and ovary in response to water deficit, heat stress, or a combination of water deficit and heat stress. Heat maps depicting the expression of transcripts encoding proteins involved in abscisic acid biosynthesis and degradation (a), the superoxide producing respiratory burst oxidase homologs (b), and auxin response factor signaling (c) are shown. Only transcripts with a significant expression level compared to control are shown. All experiments were conducted in 3 biological repeats each with tissues pooled from 8-20 different plants (depending on tissue type). Significant changes in transcript expression compared to control were defined as adjusted P < 0.05 (negative binomial Wald test followed by Benjamini–Hochberg correction). Abbreviations: WD, water deficit; HS, heat stress; WD+HS, a combination of WD and HS; RBOH, respiratory burst oxidase homolog; ABA, abscisic acid.

Analysis of the expression pattern of transcripts encoding the superoxide radical (ROS)- generating enzyme respiratory burst oxidize homolog (RBOH), that plays a key role in the regulation of ROS signaling during plant development and abiotic/biotic stresses (Suzuki et al., 2011; Wang et al., 2020; Devireddy et al., 2021; Mittler et al., 2022), in the different tissues in response to the different stress conditions further revealed that the expression of transcripts encoding numerous RBOHs is suppressed in many of the tissues in response to WD+HS (Figure 6b). The expression of transcripts encoding specific RBOH isozymes was however elevated in a few of the tissues. The complex pattern of RBOH expression in the different tissues in response to the different stress treatments could suggest that ROS levels in the different flower organs, pod and leaf are determined by an interplay between the expression of different ROS scavenging (Figure 5b) and ROS producing (Figure 6b) enzymes during the different stresses, and that this interplay is different among the different tissues.

As an example for a unique response that appeared in only one tissue type for only one stress condition, we focused on the Auxin Response Factor (ARF) pathway. As shown in Figure 6c, the expression of many transcripts regulated by the ARF pathway (GO annotation: GO:0032012; regulation of ARF protein signal transduction) was suppressed in anther specifically during a combination of WD+HS. As the expression of several ROS scavenging transcripts was significantly elevated in anther during WD+HS combination (Figure 5b), and auxin and ROS signaling interact (Blomster et al., 2011; Devireddy et al., 2021), it is possible that this unique response of the ARF pathway during WD+HS is associated with ROS and auxin responses in anther during stress combination (Figures 6c and 5b).

Taken together, the results presented in Figures 4-6 reveal that the responses of many of the different plant tissues to WD, HS, or WD+HS are mediated by different pathways, and/or different transcripts that belong to the same pathway. These could involve an interplay between and within the HSF and UPR pathways (Figure 4), the ROS scavenging and production pathways (Figures 5b and 6b), the ABA synthesis and degradation pathways (Figures 5a and 6a), and the auxin and ROS pathways (Figures 5b and 6c). Future mining efforts of the reference dataset generated by this study, for the involvement of additional pathways in the different tissues in response to the different stress conditions, could eventually lead to a more comprehensive understanding of the acclimation process of different reproductive tissues to WD, HS, and/or WD+HS. This could in turn lead to the development of new strategies to enhance the resilience of soybean to various climate change-associated stresses/stress combinations.

### Network analysis of HSFA2 in ovule in response to WD, HS, or a combination of WD+HS

To demonstrate the utility of the comparative reference transcriptomic dataset obtained by our study, we conducted a limited network regulatory study of one gene (*i*.*e*., HSFA2) using the GENIE3 platform (Huynh-Thu et al., 2010). HSFA2 is a central regulator of the HSF network in different plants (Liu and Charng, 2013; Lämke et al., 2016; Ohama et al., 2017), and is encoded in soybean by 5 genes (Figures 4a and 7). As not much is known about the response of ovary to different stresses in soybean, we focused on the response of ovary to WD, HS, or WD+HS. As shown in Figure 7a, the expression of all 5 transcripts encoding HSFA2 was unaltered in response to WD (shown by yellow triangles in Figure 7a), resulting in a very limited response of transcripts associated with this transcription factor (TF). The few HSFA2-associated transcripts that did show altered expression in Figure 7a could be regulated by other stress response regulatory transcripts that may belong to the HSF network, or to other HS/WD/WD+HS-response networks (Figures 4- 6). In contrast to WD, four HSFA2 transcripts were upregulated in ovary in response to HS (shown by red triangles in Figure 7b) and the expression of many more transcripts associated with HSFA2s was altered (Figure 7a, 7b). These included 95 transcripts that were upregulated and 84 that were downregulated. Included in these transcripts were several small heat shock proteins (sHSPs) like sHSP20 and sHSP17.6, HSFA6B, and several proteins involved in PCD regulation (Supplementary Tables S34-S36). In response to WD+HS all 5 transcripts encoding HSFA2 were upregulated (shown by red triangles in Figure 7c) resulting in an even more extensive response that included 137 and 133 transcripts that were up or down regulated, respectively, and included different sets of heat and drought response proteins, such as those encoding dehydration responsive element-binding 2C (DREB2C), and proteins involved in proline, pyruvate and phosphoenolpyruvate metabolism/signaling (Figure 7a, 7c; Supplementary Tables S34-S36). Interestingly, very little overlap was found between the transcripts associated with HSFA2 expression during WD, HS, or WD+HS in ovary (Figure 7d).

**Figure 7.**
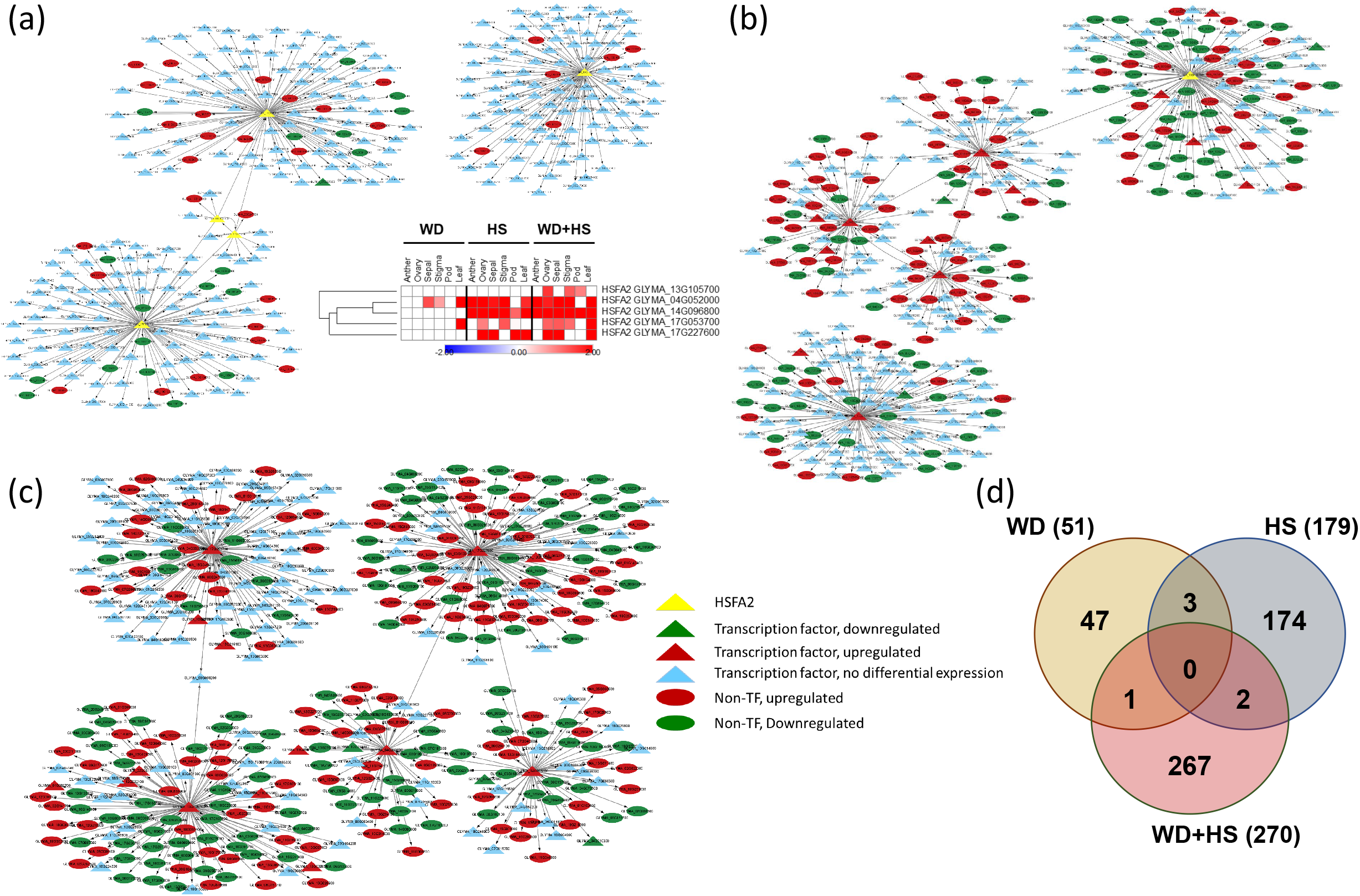
Transcriptional regulatory network analysis for heat shock transcription factor A2 (HSFA2) in ovary of soybean plants subjected to water deficit, heat stress, or a combination of water deficit and heat stress. (a) to (c), gene regulatory network maps for all five soybean HSFA2s in response to water deficit (a), heat stress (b) or water deficit and heat stress (c). A heat map for the expression of all HSFA2s in all tissues under the different conditions, extracted from Figure 4a, is included in (a). (d) Venn diagram showing the overlap between all transcripts associated with HSFA2 function under the different stress conditions (water deficit, heat stress or a combination of water deficit and heat stress). GENIE3, which infers a weighted adjacency matrix derived from Random forests using gene expression data, was used to identify HSFA2 targets, and Cytoscape3.9.1 was used to generate the different regulatory maps. A cutoff of 0.31, 0.32, 0.35 weights was used to obtain the top regulatory connections. Abbreviations: WD, water deficit; HS, heat stress, WD+HS, a combination of WD and HS; HSFA2, heat shock transcription factor A2.

Taken together, the findings presented in Figure 7 reveal that different HSFA2-associated regulatory networks are triggered in ovary during responses to WD, HS, or HS+WD, and that some transcripts belonging to these networks could also be associated with the expression of other transcripts that are unknown at present. Identifying additional HS, WD, and WD+HS networks and regulators in ovary could provide new lead pathways and genes that may be useful for breeding and engineering efforts to enhance the resilience of reproductive tissues of crops to WD, HS, or WD+HS conditions.

### Identification of reproductive tissue-specific transcripts in soybean

A key outcome of the current study is the finding that different tissues of soybean responded differently to WD, HS, and WD+HS (Figures 3-7). Augmenting the resilience of crops to different stress conditions might therefore require simultaneously altering the expression of specific pathways and/or transcripts in a tissue-specific manner. However, very few tissue-specific promoters that could be used for such efforts are available in soybean. To begin identifying tissue-specific promoters in soybean, with a focus on reproductive tissues, we identified transcripts that may be driven by these promoters based on their expression pattern under non-stress conditions in our dataset (FPKM > 2 for a specific tissue while FPKM < 1 for all other tissues). Next, we determined the differential expression of the selected transcripts in response to WD, HS, or WD+HS and kept only transcripts that were not suppressed under any of these conditions (to establish that the tissue specific transcripts identified are not suppressed during stress and their promoters could potentially be used to drive tissue- or stress-response transgenes under control conditions as well as during stress). The expression of the transcripts selected as described above was also tested in all other tissues in response to WD, HS, or WD+HS, and transcripts that responded in any other tissue to these stresses, were removed (to eliminate transcripts that are not tissue-specific under stress). Finally, the expression of the remaining reproductive tissue-specific transcripts was determined in the soybean eFP browser (http://bar.utoronto.ca/eplant_soybean/), and only transcripts that were flower- or pod-specific in both the eFP browser and our dataset were maintained. As shown in Table 1 (most abundant 5 transcripts for each tissue) and Supplemental Table S37 (full list), several reproductive tissue-specific transcripts were identified using this protocol for ovary, anther, pod, and sepal (but not stigma that had a high overlap with ovary; Figure 1b). Because the expression level of these transcripts could be regulated at the transcriptional and/or post-transcriptional level, additional studies using promoters fused to reporter genes such as green fluorescent protein are needed to determine whether the promoters driving the expression of these reproductive tissue-specific transcripts could be used in future studies as tissue-specific promoters.

**Table 1.**
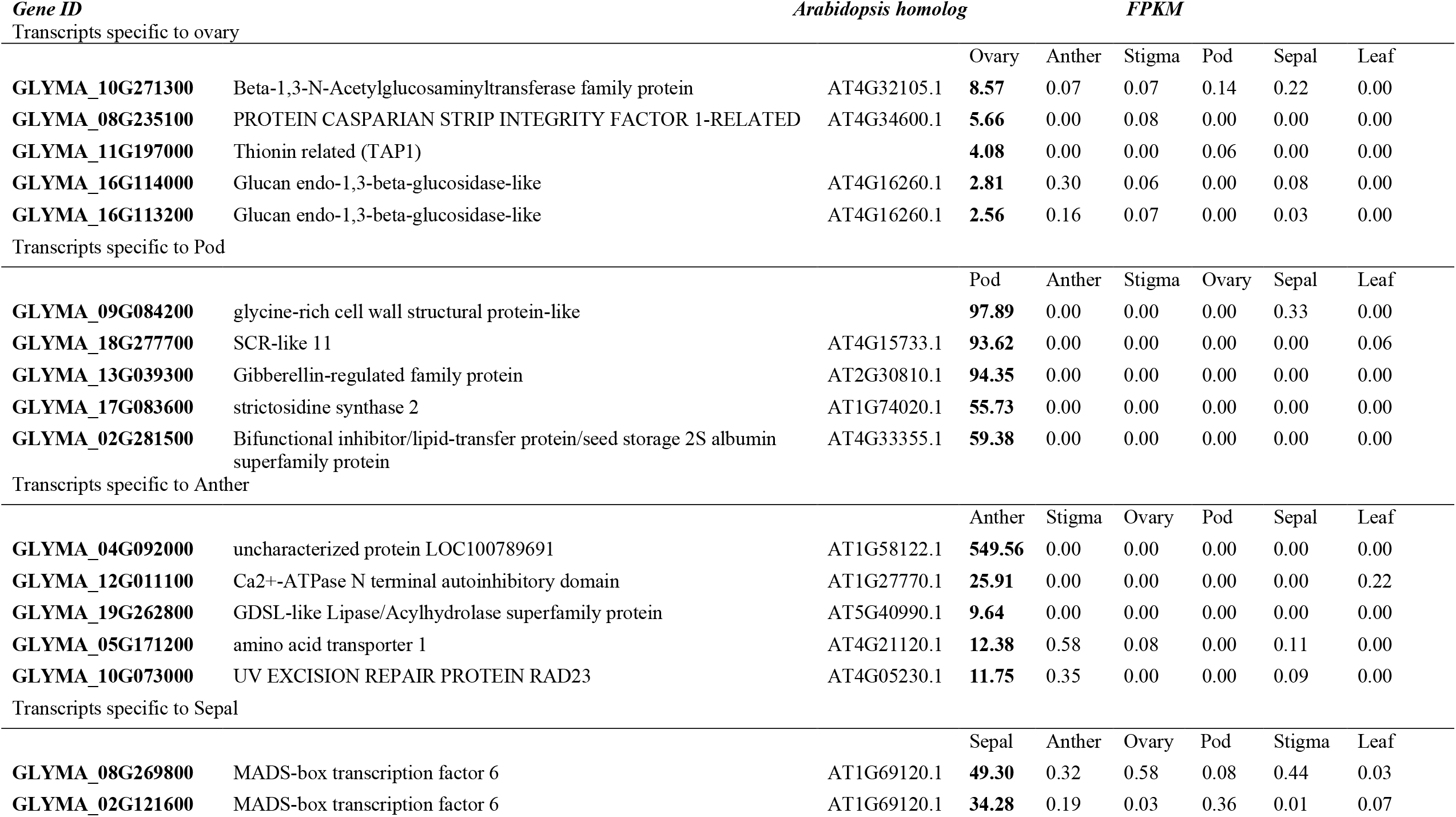
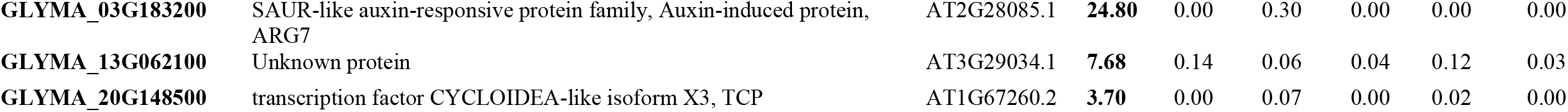
Soybean reproductive tissue-specific transcripts

## SUMMARY AND CONCLUSIONS

The comparative RNA-Seq dataset generated as a resource by this work provides an important transcriptomic reference for the expression of WD-, HS-, and/or WD+HS-response transcripts in different reproductive and vegetative tissues of soybean. Importantly, it reveals that different tissues respond differently to each of these stresses, as well as to their combination (Figures 2-7). This finding is important as it suggests that attempting to enhance the resilience of crops to different stresses and their combination might require a coordinated approach that simultaneously alters the expression of different groups of transcripts in different tissues in a stress-specific manner. In future studies the baseline reference transcriptomic dataset generated by this study could be refined and augmented using approaches such as single-cell sequencing to generate a comprehensive spatial and temporal expression atlas of the responses of each cell in each tissue to the different stressful conditions. In addition, hormone, transcript, protein, and metabolite levels in each tissue could be determined at different times during the application of WD, HS, or WD+HS to plants, and additional molecular and metabolic studies of reproductive tissues under field conditions could be conducted. Further studies focusing on many of the tissue-specific transcripts identified by this study (Table 1; Supplemental Table S37) could also identify reproductive tissue-specific promoters that could be used in future efforts to augment the tolerance of crops such as soybean to climate change-driven stresses and weather events.

## MATERIALS AND METHODS

### Soybean growth and stress treatments

Soybean (*Glycine max*, cv *Magellan*) seeds were coated with *Bradyrhizobium japonicum* inoculum (N-DURE, Verdesian Life Sciences, NC, USA) and germinated for a week in Promix BX (Premier Tech Horticulture; PA, USA), under short day growth condition (12-h light/12-h dark), at 28/24 ℃ day/night temperature and 1000 μmol photons m^-2^ s^-1^ in a growth chamber (BDR16, Conviron; Canada). The temperature of the chambers was ramped from 24 to 28 °C between 6.00-8.00 AM and decreased to 24 °C from 16.00-20.00 PM. Five-day-old seedlings were transplanted into pots containing 1 kg mixture of Promix BX and perlite (Miracle-Gro® Perlite, Miracle-Gro, Marysville, OH, USA) mixed in ratio of 10:1 and soaked with 1 l of water-fertilizer (Zack’s classic blossom booster 10-30-20; JR peters Inc., USA) mix (Sinha et al., 2022b). For the next 16-18 days, until first open flower (R1 developmental stage; Fehr et al., 1971), plants were grown under 28/24 ℃ day/night temperatures and 1000 μmol photons m^-2^ s^-1^ light intensity (12-h light/12-h dark photoperiod). Plants were irrigated twice a week with fertilizer (Zack’s classic blossom booster 10-30-20; JR peters Inc., USA; Cohen et al., 2021a; Sinha et al., 2022b). At R1, plants were randomly divided into control (CT), and 3 stress categories as water-deficit (WD), heat stress (HS), and a combination of water-deficit and heat stress (WD+HS) in four identical BDR16 growth chambers placed side-by-side in the same room (Sinha et al., 2022b). The relative humidity of chambers was maintained at about 50-60% in all chambers. The plants under WD and WD+HS treatments were irrigated daily with only 30% of the water (and fertilizer) available for transpiration as described previously (Cohen et al., 2021a; Sinha et al., 2022b), while plants in the CT and HS treatments were irrigated with 100% of the water available for transpiration. For HS and WD+HS treatments, temperature of chambers was increased to 38 °C day and 28 °C night temperature by ramping the temperature up between 6.00-8.00 AM and decreasing it down to 28 °C between 16.00-20.00 PM.

### Sample collection and RNA isolation

Sampling of all tissues begun 10-15 days after the different stress conditions were initiated (Sinha et al., 2022b; Sinha et al., 2022a; Cohen et al., 2021a). All flowers used for the transcriptomic analysis presented in this study were at stages II and III (unopen flowers undergoing self-pollination) from plants at the R2 stage (Sinha et al., 2022b). Unopened flowers from plants grown under the different growth conditions were rapidly dissected (Figure 1a) and sepal, anther, stigma, and ovary were quickly frozen in liquid nitrogen. Flower organs, leaves, and pods of soybean plants were collected between 11.30 AM-12:30 PM (Cohen et al., 2021a; Sinha et al., 2022b). All flowers used in this study were at stages II and III (unopen flowers undergoing self-pollination) from plants at the R2 stage, and all pods were at a length of about 3 cm and contained developing seeds (Sinha et al., 2022a, 2022b). For each biological repeat, flower organs were pooled together from 15-20 different plants and pods and leaves were pooled from 8-10 different plants (Sinha et al., 2022a; Cohen et al., 2021a; Sinha et al., 2022b). RNA from sepal, pod, leaf, and ovary was isolated using RNeasy plant mini kit (Qiagen, Germantown, MD, USA) whereas RNA from anther and stigma was isolated using RNeasy Micro Kit (Qiagen, Germantown, MD, USA).

### RNA sequencing and data analysis

RNA libraries were prepared using standard Illumina protocols and RNA sequencing was performed using NovaSeq 6000 PE150 by Novogene co. Ltd (https://en.novogene.com/; Sacramento, CA, USA). Read quality control was performed using Trim Galore v0.6.4 (https://www.bioinformatics.babraham.ac.uk/projects/trim_galore/) & FastQC v0.11.9 (https://www.bioinformatics.babraham.ac.uk/projects/fastqc/). The RNA-seq reads were aligned to the reference genome for Soybean - Glycine max v2.1 (downloaded from ftp://ftp.ensemblgenomes.org/pub/plants/release-51/fasta/glycine_max/dna/), using Hisat2 short read aligner (Kim et al., 2019). Intermediate file processing of sam to sorted bam conversion was carried out using samtools v1.9 (Danecek et al., 2021). Transcript abundance in levels expressed as FPKM was generated using the Cufflinks tool from the Tuxedo suite (Trapnell et al., 2012), guided by genome annotation files downloaded from the same source. Differential gene expression analysis was performed using Cuffdiff tool (Trapnell et al., 2013), from the same Tuxedo suite. Differentially expressed transcripts were defined as those that had adjusted p < 0.05 (negative binomial Wald test followed by Benjamini–Hochberg correction). Functional annotation and quantification of overrepresented GO terms (p < 0.05) were conducted using g:profiler (Raudvere et al., 2019). Venn diagrams were created in VENNY 2.1 (BioinfoGP,CNB-CSIC) and Upset Plots were created in upsetr (gehlenborglab.shinyapps.io; Conway et al., 2017). Heatmaps were generated in Morpheus (https://software.broadinstitute.org/morpheus).

### Gene Regulatory Network Analysis

The R package GENIE3 (1.18.0; Huynh-Thu et al., 2010), which infers a gene regulatory network (in the form of a weighted adjacency matrix) using gene expression data, was used to identify the targets of HSFA2 in ovary tissue using the Random Forest tree calculation method. The higher the weights, the more likely are the regulatory connections between the TF’s and their targets. A cutoff of 0.31, 0.32, 0.35 weights was used for ovary under WD, HS, and WD+HS respectively to get the top regulatory connections. This cutoff weighted adjacency matrix was later used to generate a gene regulatory network using Cytoscape tool (3.9.1; Otasek et al., 2019), an open-source platform for visualizing complex networks. For each network, a styling table was created, with up-regulated genes styled as red if the log2 fold change is greater than 0, and down-regulated genes styled as green if the log2 fold change is less than 0. In each network, transcription factor encoding transcripts were represented in a triangle shape and differentially expressed transcripts in ellipse shape. TFs are represented by blue color, whereas HSFA2 is represented by yellow color for easier identification.

### Selection of reproductive tissue-specific transcripts

Transcripts expressed in specific reproductive organs were initially identified based on FPKM levels of each transcript under nonstress (CT) conditions. Transcripts with an FPKM level of > 2 for a specific tissue (*e*.*g*., anther) and <1 for all other tissues (*e*.*g*., sepal, stigma, ovary, pod, and leaf) were selected. In the second step, differential expression of the selected transcripts was determined under WD, HS or WD+HS in the RNAseq data of each specific tissue. Transcripts with down-regulated expression were removed from the selected list (to avoid suppression of the tissue specific expression during stress). Further, the expression of transcripts selected in the second step was determined in the RNAseq datasets of all other tissues and transcripts having any differential (up/down-regulated) expression under any of the stress conditions (WD, HS, or WD+HS) in other tissues were removed. In the final step, the selected transcripts were checked for tissue specific expression in the soybean eFP browser (http://bar.utoronto.ca/eplant_soybean/) and transcripts having expression only in flowers (anther, sepal, ovary, stigma) or pods were maintained as tissue specific transcripts (Table 1; Supplemental Table S37). For the tissue-specific expression of different transcripts under control conditions (Figure 1b), the transcript expression cutoff used was > 5 FPKM.

### Statistical analysis

All experiments were conducted in 3 biological repeats each with tissues pooled from 8-20 different plants (Flower organs were pooled together from 15-20 different plants and pods and leaves were pooled from 8-10 different plants). Significant changes in transcript expression compared to control was defined as adjusted P < 0.05 (negative binomial Wald test followed by Benjamini–Hochberg correction).

## SUPPLEMENTARY TABLES

**Table S1**. List of transcripts with FPKM>5 in anther of soybean plants when grown under control (CT) condition (Figure 1b).

**Table S2**. List of transcripts with FPKM>5 in ovary of soybean plants when grown under control (CT) condition (Figure 1b).

**Table S3**. List of transcripts with FPKM>5 in sepal of soybean plants when grown under control (CT) condition (Figure 1b).

**Table S4**. List of transcripts with FPKM>5 in stigma of soybean plants when grown under control (CT) condition (Figure 1b).

**Table S5**. List of transcripts with FPKM>5 in pod of soybean plants when grown under control (CT) condition (Figure 1b).

**Table S6**. List of transcripts with FPKM>5 in leaf of soybean plants when grown under control (CT) condition (Figure 1b).

**Table S7**. List of transcripts with FPKM>5 unique to leaf of soybean plants when grown under control (CT) condition (Figure 1b).

**Table S8**. List of transcripts with FPKM>5 unique to anther of soybean plants when grown under control (CT) condition (Figure 1b).

**Table S9**. List of transcripts with FPKM>5 unique to ovary of soybean plants when grown under control (CT) condition (Figure 1b).

**Table S10**. List of transcripts with FPKM>5 unique to sepal of soybean plants when grown under control (CT) condition (Figure 1b).

**Table S11**. List of transcripts with FPKM>5 unique to pod of soybean plants when grown under control (CT) condition (Figure 1b).

**Table S12**. List of transcripts with FPKM>5 unique to stigma of soybean plants when grown under control (CT) condition (Figure 1b).

**Table S13**. List of transcripts with FPKM>5 commonly expressed in flower organs, leaf and pod of soybean plants when grown under control (CT) condition (Figure 1b).

**Table S14**. List of transcripts with FPKM>5 commonly expressed in reproductive organs of soybean plants when grown under control (CT) condition (Figure 1b).

**Table S15**. List of transcripts with FPKM>5 common between stigma and ovary of soybean plants when under control (CT) condition (Figure 1b).

**Table S16**. Transcripts differentially expressed in anther of soybean plants subjected to water deficit stress (WD).

**Table S17**. Transcripts differentially expressed in anther of soybean plants subjected to heat stress (HS).

**Table S18**. Transcripts differentially expressed in anther of soybean plants subjected to combination of water deficit and heat stress (WD+HS).

**Table S19**. Transcripts differentially expressed in ovary of soybean plants subjected to water deficit stress (WD).

**Table S20**. Transcripts differentially expressed in ovary of soybean plants subjected to heat stress (HS).

**Table S21**. Transcripts differentially expressed in ovary of soybean plants subjected to combination of water deficit and heat stress (WD+HS).

**Table S22**. Transcripts differentially expressed in sepal of soybean plants subjected to water deficit stress (WD).

**Table S23**. Transcripts differentially expressed in sepal of soybean plants subjected to heat stress (HS).

**Table S24**. Transcripts differentially expressed in sepal of soybean plants subjected to combination of water deficit and heat stress (WD+HS).

**Table S25**. Transcripts differentially expressed in stigma of soybean plants subjected to water deficit stress (WD).

**Table S26**. Transcripts differentially expressed in stigma of soybean plants subjected to heat stress (HS).

**Table S27**. Transcripts differentially expressed in stigma of soybean plants subjected to combination of water deficit and heat stress (WD+HS).

**Table S28**. Transcripts differentially expressed in pod of soybean plants subjected to water deficit stress (WD).

**Table S29**. Transcripts differentially expressed in pod of soybean plants subjected to heat stress (HS).

**Table S30**. Transcripts differentially expressed in pod of soybean plants subjected to combination of water deficit and heat stress (WD+HS).

**Table S31**. Transcripts differentially expressed in leaf of soybean plants subjected to water deficit stress (WD).

**Table S32**. Transcripts differentially expressed in leaf of soybean plants subjected to heat stress (HS).

**Table S33**. Transcripts differentially expressed in leaf of soybean plants subjected to combination of water deficit and heat stress (WD+HS).

**Table S34**. List of transcripts associated with HSFA2 transcription factor in ovule of soybean plants subjected to WD stress.

**Table S35**. List of transcripts associated with HSFA2 transcription factor in ovule of soybean plants subjected to HS stress.

**Table S36**. List of transcripts associated with HSFA2 transcription factor in ovule of soybean plants subjected to WD+HS stress.

**Table S37**. Soybean reproductive tissue-specific transcripts.

## ACKNOWLEDGMENTS

This work was supported by funding from the National Science Foundation (IOS-2110017, IOS- 1353886, IOS-1932639), Interdisciplinary Plant Group, and University of Missouri.

## AUTHOR CONTRIBUTIONS

R.S., S.P.I, M.A.P.V, A.T., and B.S. performed experiments and analyzed the data. R.M., F.B.F, R.S., T.J., M.A.P.V and S.I.Z. designed experiments, analyzed the data, and/or wrote the manuscript.

## DATA AVAILABILITY

Transcript abundance and differentially expressed transcripts can be accessed interactively via the Differential Expression tool in SoyKB (https://soykb.org/DiffExp/diffExp.php; Joshi et al., 2012, 2014), a comprehensive all-inclusive web resource for soybean. RNA-Seq data was deposited in Gene Expression Omnibus (GEO), under the following accession numbers: GSE218146, GSE213479, GSE218146, and GSE186317.

